# Harmonization of IGF1 immunoassay methods using an LC-MS/MS method and associated normative dataset

**DOI:** 10.64898/2026.02.15.706059

**Authors:** E.G.W.M Lentjes, M.S Pratt, I.P Kema, M van Faassen, R Musson, M.J. Vos

## Abstract

**Objective:** Generation and testing of IGF1 reference materials (RM), suitable for the harmonization of immunoassay (IA) and LC-MS/MS methods for the IGF1 determination in blood. In addition, establishment of age related reference intervals for men and women.

**Methods:** In a split sample study of 42 patients, and 30 healthy volunteers we tested the commutability of four RMs for IGF1, using four commercial IAs and an LC-MS/MS method. A new set of age dependent reference intervals was established using Lifelines biobank samples, based on the IGF1 LC-MS/MS method.

**Results:** The four RMs were found to be commutable, except the RM with the lowest concentration measured with the Siemens Immulite method. The value assignment of the RMs was based on the IGF1 LC-MS/MS method, which was calibrated against WHO international standard 02/254. LC-MS/MS results were on average about 0 to 60% lower than those of the immunoassays. Combining the recalculated IGF1 results in patient samples from a former study with the data from healthy volunteers in this study, showed a reduction in the variation of the data points (standard error of estimate) of 42% and 62% respectively.

**Conclusion:** Commutable RMs for IGF1 can be made from serum of healthy blood donors. However, it remains necessary to test the commutability of these RMs in IAs that were not included in this study. By harmonizing methods using the four RMs, the same age-related reference intervals can be used.

## Introduction

Insulin-like Growth Factor 1 (IGF1) is a 7.5 kDa protein that plays a crucial role in cell growth, tissue repair, and metabolism. It is primarily produced in the liver in response to growth hormone and circulates in the blood bound to specific transport proteins, mainly IGF-BP3 and ALS (Acid Labile Substance). IGF1 is essential during childhood for normal growth and development. In adults, it supports muscle repair, bone density, and metabolic processes. IGF1 is also used in anti-doping testing, together with procollagen III amino-terminal propeptide, to detect GH use in sports (1) (2).

Although all current IGF1 assay methods are calibrated against the same WHO international standard (WHO IS 02/254), differences in results still exist between various immunoassays. In addition, plasma IGF1 concentrations vary significantly with age, but other factors like protein intake, stress, intense exercise, several hormones, either endogenous or taken as a drug, also may influence IGF1 levels (3). For a recent review about (patho)physiological and analytical issues concerning IGF1 levels see Huang et al (4). Another source of confusion is the variety of method-dependent reference intervals currently in use. Collectively, these variations lead to divergent clinical decisions regarding patient treatment depending on the immunoassay employed (5)(6)(7)(8), underscoring the need for harmonized methods and robust, age-specific reference intervals based on an extensive normative dataset. Such rigorous intervals, however, may not be easily interchangeable across methods due to substantial inter-assay variability (9), rendering the use of the same reference dataset across different IAs unsuitable.

Standardization of these methods is not yet possible due to the absence of commutable reference material and a reference method. An alternative approach is the harmonization of methods using commutable RM, intended for use as a calibrator. In 2013, the Endocrinology section of the Dutch EQAS SKML (Foundation for Quality Assessment in Laboratory Medicine; SKML) initiated a project in the Netherlands to harmonize IGF1 methods, ensuring that measurement results are independent of the assay used.

Already in 2004, a commutable serum growth hormone RM was developed in the Netherlands, which led to a reduction in the between-laboratory coefficient of variation (CV) and has been successfully used for years (10). This same sample was also found to be commutable for IGF1 methods and has been in use for several years. However, this single RM proved less effective in reducing the between-laboratory CV for IGF1 compared to GH. Therefore, four new RMs were developed to span a broader concentration range and to mitigate matrix effects specific to IGF1, ultimately aiming to standardize IGF1 measurements and improve diagnostic reliability. In addition, a successful harmonization of methods enables the introduction of universal reference intervals that apply to all assays. This prevents the need for each laboratory to determine its own reference intervals, resulting in significant cost savings and ultimately promotes more uniform clinical decisions.

As part of this effort, IGF1 concentrations were measured in a large reference population using a validated LC-MS/MS method to serve as a calibration anchor. This enabled the creation of harmonized calibration curves for each assay and provided a robust foundation for establishing age- and sex-specific reference intervals.

## Materials and methods

### Samples

#### RM samples

RM samples were prepared from the serum of blood bank donors. Donors were selected based on age to obtain variations in IGF1 concentrations. To ensure a sufficient number of aliquots per concentration, a maximum of two donations were pooled. The 400 µL aliquots are stored at minus 80°C. Four potential calibrators were produced, with IGF1concentrations ranging from 7 to 32 nmol/L, as determined by LC-MS/MS.

#### Patient samples

Patient samples (from adults and children) were collected in three hospital laboratories using leftover samples from IGF1 measurements, provided no objection was raised. Exclusion criteria included conditions affecting growth, liver and kidney dysfunction and the use of chemotherapeutic agents. The materials were anonymized, aliquoted, and frozen at -20°C until shipment.

For the harmonization and commutability study, 42 patient samples were used. These samples were aliquoted and sent frozen to various laboratories.

#### Samples for reference intervals

In a previous study, reference intervals for IGF1 were established using the Diasorin Liaison assay. These included 698 men and 901 women (28% aged 0–18 years and 72% aged >18 years). The samples consisted of anonymized leftover patient samples used for screening purposes (classified as non-WMO research, thus exempt from Medical Ethical Committee approval) or were samples from healthy individuals in control groups of studies (with informed consent). The samples for these reference intervals were collected in two laboratories, and IGF1 was measured. Each run included a harmonization sample (the harmonization sample created in 2004, as mentioned in the introduction). Since this harmonization sample was also measured using LC-MS/MS, the original IGF1 results could be recalculated to align with the LC-MS/MS method.

For the establishment of LC-MS/MS (11) based reference intervals, samples from healthy volunteers (760 males and 760 females; 58% 8 to 18 yr and 42% >18yr) were obtained from the Lifelines biobank (12) (www.lifelines.nl; Groningen, the Netherlands). Lifelines is a prospective population-based cohort study of 167,729 persons in the North of the Netherlands, using a unique three-generation design with a focus on multi-morbidity and complex genetics.

The age range of the donors was from 8 to 94 years and were from apparently healthy volunteers. Samples were aliquoted at the Lifelines site and sent to the UMC Utrecht on dry ice. The aliquots were kept frozen at -80°C and subsequently shipped from the UMC Utrecht to University Medical Center Groningen on dry ice.

## Methods

The IGF1 analysis methods currently in use in the Netherlands were applied in this study. Analyses were performed according to the manufacturers’ specifications using the Liaison (DiaSorin), Cobas (Roche Diagnostics), iSYS (IDS), and Immulite 2000 (Siemens Healthineers) platforms. All methods were calibrated against the WHO international standard (IS) 02/254 (13).

All laboratories were instructed to perform calibration before analysis to ensure that control measurements were validated before proceeding with sample testing.

### Reference intervals

For the establishment of LC-MS/MS based reference intervals, all 1520 samples obtained from the Lifelines Biobank were also analyzed using LC-MS/MS which was also calibrated against WHO IS 02/254 (11). To enhance the coverage of IGF-1 reference intervals in early infancy as well as adulthood, an additional 1599 datapoints were added as described above.

To determine the reference intervals, the LMS method was used, first developed by Cole (14) and later extended by Cole and Green (15). The L, M, and S parameters were calculated for each year of life, allowing the Z-score of an IGF1 result to be determined using the formula: SDS = [(IGF1 result/M)^L^ - 1] /[L x S]. LMS Chartmaker was used for this analysis (16).

### Statistical analysis

The IFCC Working Group on Commutability has published a procedure in which the bias of an RM is compared to that of clinical samples (CS) against a predefined criterion based on medically relevant differences (17). We used the Excel sheets provided in the data supplements of the online version of this article for our calculations. Briefly: in experiment 1, the four calibrators were measured alongside patient samples and in experiment 2 the RMs were measured together with samples from healthy volunteers (from Lifelines Biobank). Measurements followed a fixed protocol and were conducted in a single run, with the RMs placed between the patient samples. Each sample was measured in duplicate. Note: experiment 1 and Experiment 2, which was conducted six months later, were carried out in entirely different laboratories.

For the Deming regression method, we used the GraphPad® Prism (GraphPad Software, San Diego, USA).

Linear regression calculations were performed using the SPSS® statistical package (version 30.0). For the reference intervals we tested the difference between the former, recalculated, data and the newly obtained data from the Lifelines samples. For this, only the data from individuals older than 8 years were used, as Lifelines samples are only collected from the age of 8 onwards. To compare the two groups, the differences in IGF1 results were calculated relative to the median of all data for men or women (derived from the LMS calculation) of the computed reference intervals. Since the distribution around this median was not normally distributed, a non-parametric Mann-Whitney U test was used.

## Results

The commutability study showed that all four RMs met the criteria for all immunoassays (IAs), set out in the IFCC Working Group Commutability procedure, except for RM1 which showed non-commutability with the Immulite assay. RM1, which had the lowest IGF1 concentration, deviated significantly for the Immulite in various comparisons with three other immunoassays and the LC-MS/MS method.

### Initial evaluation of immunoassays versus LC-MS/MS

In the first experiment, the RMs and patient samples were measured using four immunoassays and the LC-MS/MS method according to the IFCC protocol. In a second experiment, samples from healthy individuals were used. The slopes and intercepts of the correlations between the IAs and the LC-MS/MS method are presented in Tables 1 and 2, respectively. Figure 1 displays the results of the IGF1 measurements in patient samples and healthy volunteers, respectively. It is noteworthy that IGF1 concentrations measured by LC-MS/MS are consistently lower than those measured by the IAs. The differences ranges from negligible up to approximately 60%, depending on the assay and concentration level.

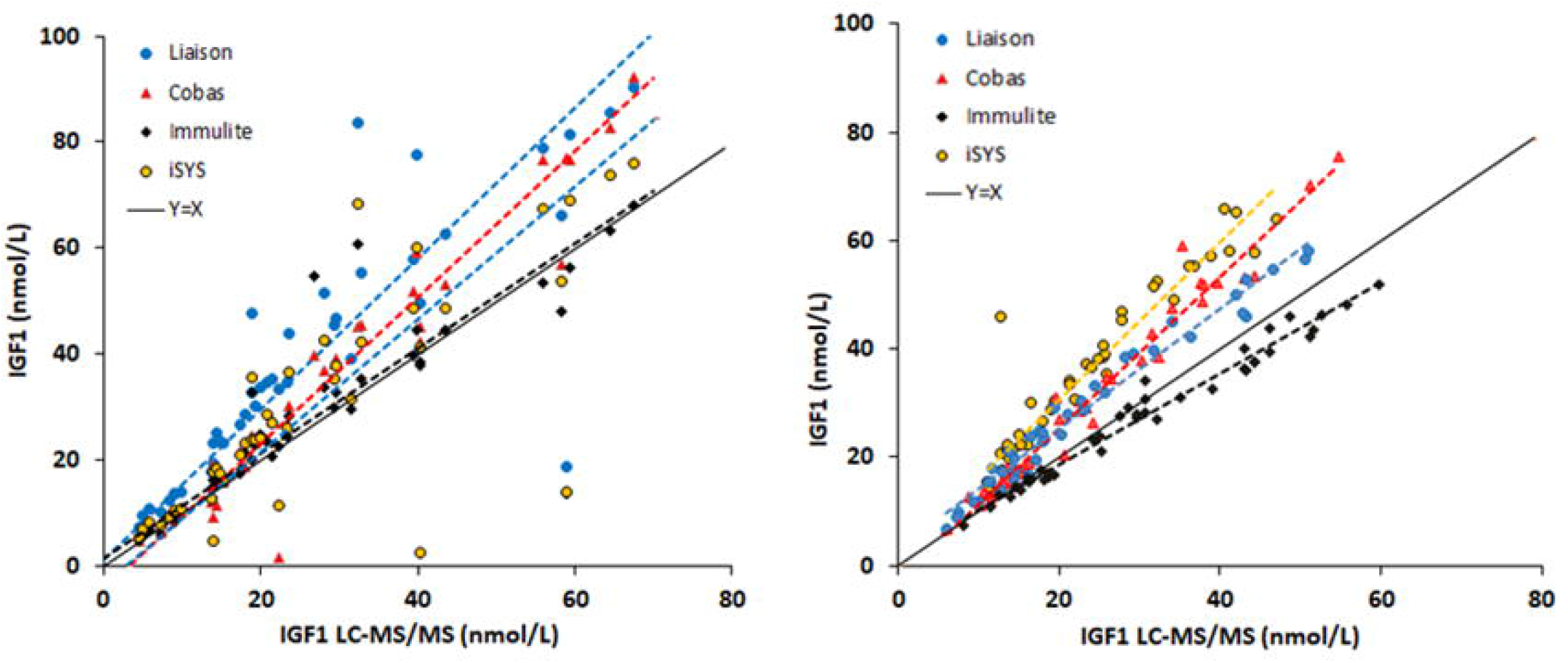

### Outliers and deviations

A striking observation in Figure 1 is the presence of 8 patient samples that deviate from the LC-MS/MS values, showing both highly increased and decreased IGF1 results when measured by IAs. These deviations are not attributable to a particular method, the analysis in a specific laboratory or to any of the three laboratories that collected the samples. Given that each sample was measured in duplicate, the values represent true analytical results rather than random error. In Figure 1, for the healthy volunteers, only one outlier is visible. Additionally, it appears that the slopes of the correlations differ (see Tables 1 and 2).

### Inter-laboratory comparison in assay performance

The RMs were measured alongside the samples from the patients and the healthy individuals. The measured IGF1 concentrations are shown in Figure 2. Notably, The Immulite method shows a distinctly different slope and intercept in two of the laboratories compared to other the three IAs.

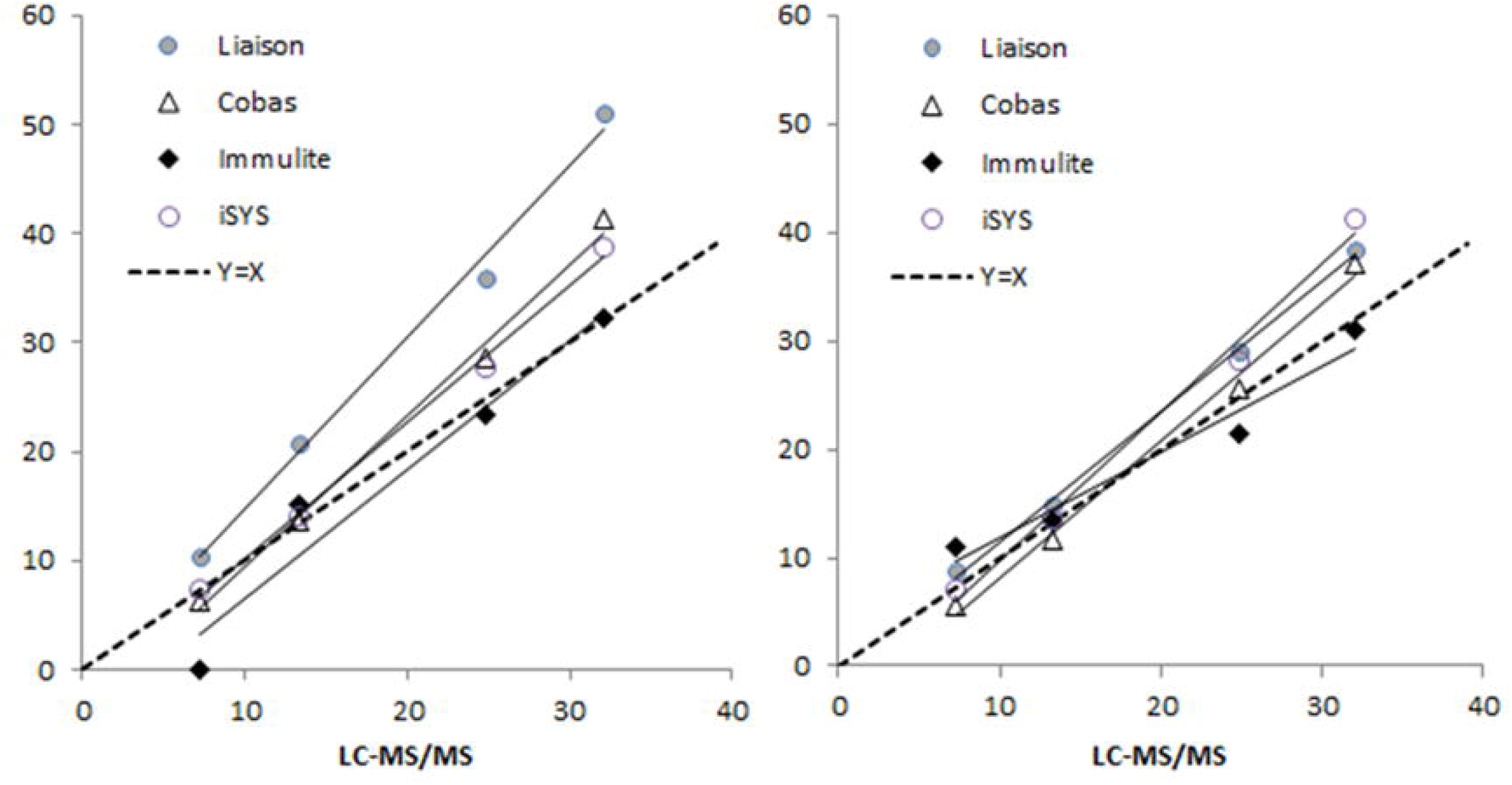

Notice that the regression lines for the Liaison and the iSYS have shifted between figures 1 and 2. This is explained by the fact that patient samples and the healthy volunteer samples were analyzed in different laboratories for both assays. Nevertheless, all measurements were performed only after thorough calibration and system verification procedures.

### Recalibration using commutable reference materials

Subsequently, the IGF1 results of both the patient samples and healthy volunteers were recalculated using the established calibration curves based on the commutable RMs: i.e. RM1-4 for the Cobas, iSYS, and Liaison, and RM2-4 for the Immulite. The recalibrated results are presented in Figure 3. The regression lines are now more parallel and closely aligned with the identity line, particularly for Cobas, iSYS, and Liaison, indicating a substantial improvement in assay harmonization across the entire concentration range.

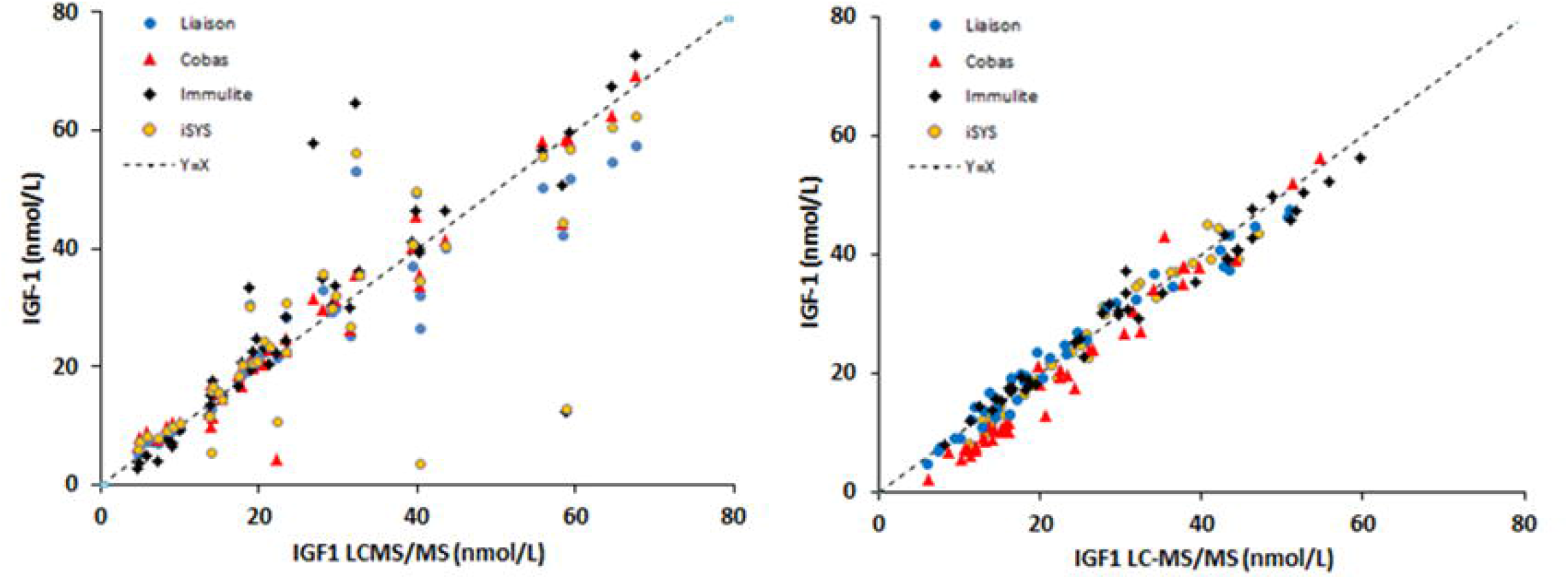

To quantify the impact of recalibration, the standard error of estimate (SEE) was calculated for both the patient and healthy groups before and after applying the RMs. As shown in Table 3, recalibration led to a substantial reduction in variability: the SEE decreased by 62% for the healthy volunteer group and by 42% for the patient group.

### Establishment of an IGF1 LC-MS/MS reference dataset

For the establishment of the reference intervals we used the LMS method. For men we had in total 1458 data points and for women 1661 (Figure 3). The -2, -1, median, +1 and +2 SD lines were calculated and plotted into the figures.

The data from the former reference study were recalculated and compared to the newly obtained values. To determine whether the two datasets were reasonably comparable, a non-parametric Mann Whitney-U test was performed on the differences in IGF1 results (of each dataset) relative to the median of the final reference intervals. No difference was observed in males (p=0.061), whereas a difference was found in females (p=0.01). The biggest difference between the groups was in the age range of 40-60 years. For the age range of 8-40 years, and above 60 years there was no difference. The table with the reference intervals by age and the corresponding LMS values can be accessed via the following link: https://doi.org/10.5281/zenodo.15799741

## Discussion

This study on the harmonization of the IGF1 methods was initiated in response to the substantial discrepancies observed between the various IGF1 measurement methods in the 2024 Dutch EQAS. The IGF1 results differed by a factor of 1,5 to 2 between the various IAs. Although frozen serum from healthy blood bank donors was used for harmonization and the various methods were all calibrated against WHO IS 02/254. In 2020, Lee et al reported also wide variations between three IA platforms and an LC-MS/MS method, all methods calibrated against WHO IS 02/254 and compared the reference ranges, obtained from the manufacturers, which also showed differences (18). Substantial inter-method differences among IGF1 assays have been reported earlier (19) (20), but these studies were mostly done with ELISAs or radiometric assays and were calibrated against WHO IS 87/518, which was found to be of low purity containing 44% IGF1 (21). Not only the methods showed large variation, but in an UK NEQAS study reported in 2007 by Pokrajac et al. the authors mentioned that age-related reference intervals varied significantly between laboratories, even when using the same method. This led to considerable differences in the interpretation of IGF1 results (9). More recently, Chanson et al showed in 2016 that when establishing IGF1 reference intervals using six immunoassays, the four methods calibrated against WHO 02/254 exhibited only moderate agreement (Table 3: agreement expressed as weighted kappa) (22). Additionally, the concordance between manufacturers’ reference intervals and those obtained in their study was generally poor. Not only is there a substantial inter-method variability, but also the reference intervals that are used by the various laboratories, show much variation. This means that calculating the SDS of the IGF1 result will not lead to better agreement between methods and laboratories. Indeed, Mavromati et al in 2017, showed in over 100 patients with growth hormone disorders marked variability in IGF-I blood concentrations and in IGF-I calculated SDS values obtained with each of the six immunoassays (23). Also, Varewijck et al, who investigated the use of two normative datasets for the calculation of IGF1 Z-scores in 102 GH deficient subjects concluded that there was a large difference in the interpretation of the results (7). This would potentially lead to misclassifications of the patients, which is clearly shown in the recent study by Postma et al., where significant analytical variation was observed in measured IGF1 concentrations (all methods calibrated against WHO IS 02/254) and in the SDS values, which was even greater. In addition, among the physicians clinical interpretation of IGF1 results in the five patient cases varied widely (5). This study highlights the need for multiple, commutable RM samples covering a broader concentration range, rather than relying on a single harmonization sample.

Additionally, there remains a responsibility for physicians to adhere closely to the guidelines. In this study, donor serum from healthy individuals was used for the RMs, with the expectation of normal IGF-binding proteins in the serum (not measured) and a high likelihood of obtaining commutable RMs. The results show that for the three IAs, Liaison, iSYS and Cobas, and the LC-MS/MS method all four RMs are commutable. However, for the Immulite, the lowest RM was found to be non-commutable. In addition, the regression line through the three RM values measured with the Immulite exhibits a significantly different slope compared to the three other immunoassays. The cause of this discrepancy is unclear but may be related to the calibration of the method, variation in antibodies and/or the method used for dissociation of IGF1 from the binding proteins. Recalculating the Immulite IGF1 values based on RM2-4 results for both patient and healthy individual samples provides a good correction. The reason for the different behavior of the Immulite assay on the RM1 sample is not clear. The serum selection for the RMs was based on age, which may have influenced the RM with the lowest concentration (e.g., differences in IGF1 binding proteins or perhaps the presence of heterophilic antibodies).

IGF1 concentrations measured on the LC-MS/MS are on average lower compared to the IAs. This was also observed earlier (24) (11) when comparing the mass spectrometry method to the iSYS IA. Reasons that may give an explanation for this are the following: specificity of the LC-MS/MS for the target IGF1 molecule, presence of IGF1 variants and degradation products, which can sometimes be missed by either the IA or LC-MS/MS, differences in the dissociation step to release IGF-1, or more interference of IAs for matrix effects (binding proteins, heterophilic antibodies, rheumatoid factors). A striking observation in Figure 1, where the RMs were measured among patient samples, is that six samples show significant discrepancies in IGF1 results. Since these were duplicate measurements with minimal variation between the two results, they appear to be real values. The discrepancies (both significantly elevated and reduced values) were observed across multiple analyzers in different laboratories. All laboratories received aliquots from the exact same samples. Unfortunately, further testing was not possible because these were leftover patient samples with limited volume. Comparable discrepancies of IA results, compared to the LC-MS/MS in individual samples have also been reported by Lee et al (18). Since the LC-MS/MS method did not show deviations compared to the other immunoassays, it is unlikely that IGF1 variants (25) were involved, as the mass difference of the variants compared to the wild type IGF1 is usually substantial. However, variable concentrations of IGF-binding proteins (26) or heterophilic antibodies in the serum (27)(28) may have contributed to the inconsistent IA results. In the figure comparing the immunoassays with LC-MS/MS in healthy individuals, only a single outlier was observed. Since the patient serum samples underwent one additional freeze-thaw cycle compared to the samples from healthy individuals (Lifelines samples), 12 aliquots of Lifelines samples (IGF1 range 6 to 48 nmol/L) were subjected to an extra freeze-thaw cycle. This resulted in 18% higher readings for the iSYS and 16% higher readings for the Liaison.

This suggests that the additional freeze–thaw cycle may have altered the sample matrix (29), for example by partially dissociating IGF-1 from its binding proteins or modifying tertiary protein structure, thereby increasing the immunoassay-detectable fraction of IGF1 or non-selective binding of IA antibodies to other proteins, without affecting its actual concentration. This effect can be assay-specific, as antibodies used and the IGF binding protein disruption step varies between manufacturers. Additionally, the spread around the regression line increased after the extra freeze-thaw cycle. The effects, however, were too limited to explain the substantial discrepancies observed in the six patient samples.

The data in Tables 1 and 2 show that even within a single method, differences exist between laboratories, despite the fact that instructions of the manufacturers were carefully followed. This is also visible in Figure 2, which shows the slope and intercept of the RMs for the four methods. It is possible that these differences are due to lot-to-lot reagent variations (30)(26).

After recalculating all IGF1 values for both patient and healthy samples based on the RMs, a significant reduction in inter-method variability was observed. This is evident in Figure 4, for both patient and healthy samples. Table 3 demonstrates the reduction in the SEE of the overall regression line and the increase in the coefficient of determination, i.e. R^2^. These findings confirm that the use of matrix-matched, commutable reference materials can significantly enhance inter-assay comparability and measurement precision.

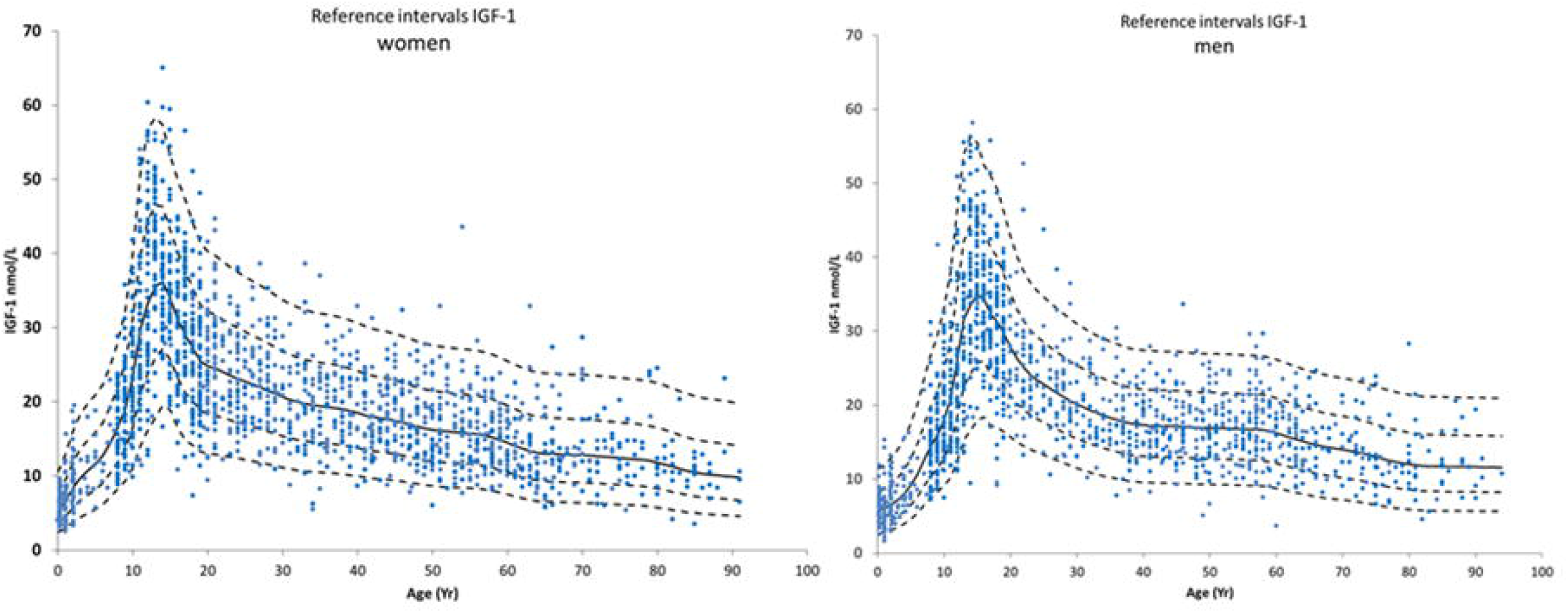

The use of commutable RMs eliminates the need to establish separate reference intervals for each commercial method, as was done by Chanson et al. (22). However, within a single method, differences between laboratories persist, despite proper maintenance and calibration. The use of commutable RMs by each laboratory helps correct these discrepancies.

We calculated reference intervals for men and women, combining the former IGF1 data, which were recalculated to the LC-MS/MS level, with the IGF1 data from the Lifelines Biobank samples. This was possible because these former data were analyzed together with a harmonization sample, which is still available, and was analyzed with the LC-MS/MS. For the male data, the recalculated data and the newly established did not show a difference in the statistical test. However, the two data groups for women did show a difference. Upon closer inspection, this was mainly caused by the age range of 40 to 60 years. The median IGF1 values of the patient group were 3.3 nmol/L higher in this range (17.27 vs. 13.97 nmol/L, and 310 vs 84 data points respectively). A possible reason is that the number of data points is much higher for the patient group, which may have resulted in a skewed distribution. Other reasons for the higher values can be insulin resistance or type 2 diabetes mellitus, or underlying medical conditions like (undetected) acromegaly, high protein or caloric intake.

It would be preferable for manufacturers to adopt these RMs for standardization, especially when reliable reference methods become available. This would enable the establishment of universal age- and sex-adjusted reference intervals with sufficient data points. This will lead to better diagnoses of growth disorders in children and adults, as well as accurate therapy and monitoring thereof. In addition it will save time and costs as well.

