## Supplementary material for "Harmonization of IGF1 immunoassay methods using an LC-MS/MS method and associated normative dataset": Figure legends

Figure 1: IGF1 results in patient samples (left) and samples from healthy individuals (right) determined with the four immunoassays, plotted against the LC-MS/MS values. The solid line indicates the line of identity (y = x).

Figure 2: IGF1 results of the RMs measured with four IAs, plotted against the LC-MS/MS values. On the left the results of experiment 1 and on the right of experiment 2, measured in four different laboratories. The dashed line indicates the line of identity (y = x).

Figure 3: IGF1 results after recalculation at RM1-4 or RM2-4 for the Immulite. On the left the results of the comparisons with patient samples and right with samples from healthy individuals. The dashed line indicates the line of identity (y = x). Note that the left and right panels reflect analyses performed in different laboratories.

Figure 4: Reference intervals for IGF-1 in women (left) and men (right). Each plot displays individual IGF-1 measurements (blue dots) as a function of age. The solid line represents the median (0 SDS), while the dashed lines indicate the -2 SDS, -1 SDS, +1 SDS, and +2 SDS reference limits.
